# A common *cis*-regulatory variant impacts normal-range and disease-associated human facial shape through regulation of *PKDCC* during chondrogenesis

**DOI:** 10.1101/2022.09.05.506587

**Authors:** Jaaved Mohammed, Neha Arora, Harold S. Matthews, Karissa Hansen, Maram Bader, John R. Shaffer, Seth M. Weinberg, Tomek Swigut, Peter Claes, Licia Selleri, Joanna Wysocka

## Abstract

Genome-wide association studies (GWAS) identified thousands of genetic variants linked to phenotypic traits and disease risk. However, mechanistic understanding of how GWAS variants influence complex morphological traits and can, in certain cases, simultaneously confer normal-range phenotypic variation and disease predisposition, is still largely lacking. Here, we focus on *rs6740960*, a single nucleotide polymorphism (SNP) at the 2p21 locus, which in GWAS studies has been associated both with normal-range variation in jaw shape and with an increased risk of non-syndromic orofacial clefting. Using *in vitro* derived embryonic cell types relevant for human facial morphogenesis, we show that this SNP resides in an enhancer that regulates chondrocytic expression of *PKDCC* - a gene encoding a tyrosine kinase involved in chondrogenesis and skeletal development. In agreement, *rs6740960* SNP is sufficient to confer a large difference in acetylation of its cognate enhancer preferentially in chondrocytes. By deploying dense landmark morphometric analysis of skull elements in mice, we show that changes in *Pkdcc* dosage are associated with quantitative changes in maxilla, mandible, and palatine bone shape that are concordant with the facial phenotypes and disease predisposition seen in humans. We further demonstrate that the frequency of the *rs6740960* variant strongly deviated among different human populations, and that the activity of its cognate enhancer diverged in hominids. Our study provides a mechanistic explanation of how a common SNP can mediate normal-range and disease-associated morphological variation, with implications for the evolution of human facial features.

## Introduction

The face is one of the most complex structures of the human body and has changed dramatically over the course of human evolution (Mitteroecker et al. 2004). In addition to facilitating coordination of multiple sensory organs, protection of the brain and a number of biological adaptations, the human craniofacial complex is a target of sexual selection and plays key roles in communication and other social interactions (Guo et al. 2014; Sheehan and Nachman 2014). The human face shape varies tremendously across individuals, much of which is genetically encoded, as manifested by the high similarity of facial features in monozygotic twins and by familial resemblances [reviewed in (Naqvi et al. 2022)]. Recently, much progress has been made in understanding the genetic underpinnings of the normal-range facial variation in human populations through Genome-Wide Association Studies (GWAS) [reviewed in (Weinberg et al. 2019; Naqvi et al. 2022)]. Collectively, these studies have uncovered over 300 loci associated with different aspects of facial shape, indicating that - perhaps not surprisingly - facial shape is a highly polygenic trait [reviewed in (Naqvi et al. 2022)]. Furthermore, as has been seen for many other human phenotypic traits, facial shape GWAS variants map to the non-coding parts of the genome and are especially enriched within *cis-*regulatory elements, such as enhancers (Claes et al. 2018). Enhancers are modular genetic elements that activate the expression of their target genes over long genomic distances and in a tissue-specific manner [reviewed in (Shlyueva et al. 2014; Heinz et al. 2015; Long et al. 2016; Furlong and Levine 2018)]. As would be expected, enrichment of facial GWAS signals in enhancers is specific to cell types relevant for facial morphogenesis during development, including early fetal facial tissue, *in vitro* derived embryonic facial progenitors called cranial neural crest cells (CNCCs) and their more differentiated descendants, such as cranial chondrocytes (Claes et al. 2018; Naqvi et al. 2021; White et al. 2021). These observations support the *cis-*regulatory origins of normal-range human facial diversity, whereby non-coding genetic variants affect enhancer function and lead to quantitative changes in gene expression in cell types contributing to facial morphogenesis during development.

In addition to the genetic underpinnings of normal-range facial variation, GWAS of non-syndromic cleft lip/palate (nsCL/P) has revealed the polygenic architecture of this common birth defect, with risk variants again mapping mostly to the non-coding parts of the genome (Leslie 2022; Naqvi et al. 2022; Weinberg 2022). While CL/P co-occurs with several Mendelian craniofacial syndromes, the majority of cases (~70%) are non-syndromic (meaning, occurring in the absence of other manifestations) and characterized by complex inheritance. We and others previously noted an overlap between sequence variants associated with nsCL/P risk and normal-range face shape variation (Boehringer et al. 2011; Indencleef et al. 2018; Indencleef et al. 2021). For example, the *rs6740960* variant at the 2p21 locus affects the normal-range jaw shape and confers nsCL/P risk in Europeans (Ludwig et al. 2017; Claes et al. 2018). Interestingly, such SNPs at the intersection of normal and dysmorphic facial features show high heterozygosity in the analyzed populations and affect diverse aspects of the normal-range facial morphology, such as shape of jaw, philtrum, or nose (Indencleef et al. 2018). These observations raise a question of how such common genetic variants simultaneously confer disease predisposition and normal-range phenotypic variation of distinct facial regions.

Despite the enormous progress in understanding genetics of the normal-range and disease-associated phenotypic variation of facial features and other human traits, functional follow-up studies have lagged behind. Such studies may involve discovery of functional variants directly relevant for gene expression, identifying the specific cell types and spatiotemporal contexts they act in, and pinpointing the genes they regulate [for example: (Sekar et al. 2016; Small et al. 2018; Caliskan et al. 2021; Sinnott-Armstrong et al. 2021)]. Even more rudimentary is our understanding of mechanisms by which changes in gene dosage associated with common genetic variation translate into complex human phenotypes. This scarcity of mechanistic discernment is not unique to the face, but emerges as a general bottleneck in translating GWAS results to genuine insights into molecular and cellular processes underlying complex traits and diseases, and utilizing them for new therapeutic strategies (Cano-Gamez and Trynka 2020).

Here, we deploy an integrated ‘SNP-to-phenotype’ approach, combining trait-relevant *in vitro* human stem cell models with *in vivo* studies in mice to probe the mechanism by which a specific GWAS-identified variant, *rs6740960*, affects both normal-range facial shape and confers nsCL/P risk. We identify a craniofacial enhancer harboring the *rs6740960* SNP, characterize its cell-type specific and facial-domain restricted activity pattern, and describe changes in activity during hominid evolution. Using *in situ* chromatin conformation capture experiments, we link the *rs6740960* cognate enhancer to its target gene, *PKDCC*. Through genome editing, we then uncover chondrocyte-specific impact of this enhancer on the *PKDCC* dosage. Finally, using micro-computed tomography (micro-CT) coupled with dense landmark morphometric analysis of skull elements, we demonstrate the phenotypic impact of *Pkdcc* dosage on mouse craniofacial development and reveal its striking concordance with the lower jaw phenotypes and clefting predisposition associated with the *rs6740960* SNP in humans.

## Results

### Common genetic variant at 2p21 associated with lower jaw shape variation and susceptibility to clefting shows large frequency shifts in human populations

Our recent multivariate GWAS studies of normal-range facial variation in two independent European ancestry populations revealed association of non-coding genetic variants at the 2p21 locus with lower jaw and chin shape (Claes et al. 2018; White et al. 2021). The lead SNP at this locus, *rs6740960* (A/T) (lowest meta-analysis p-value = 3.39 × 10^−36^), was significantly associated with shape variation over the entire face, however, the most pronounced effects were seen in the lower face, especially in the jaw region (**Figure 1A, Figure 1- Figure Supplement 1**). Specifically, the “T” allele (Allele Frequency = 0.497 (Claes et al. 2018)) is associated with an outward protrusion of the lower jaw and zygomatic region, coupled with retrusion of the entire central midface. Interestingly, this same lead SNP was significantly associated with non-syndromic cleft-lip and palate (nsCL/P) in a separate GWAS from an independent cohort of European ancestry individuals (Ludwig et al. 2017). The risk allele for nsCL/P was the “T” allele (p-value = 5.71 × 10^−13^) (**Figure 1A**).

**Figure 1:**
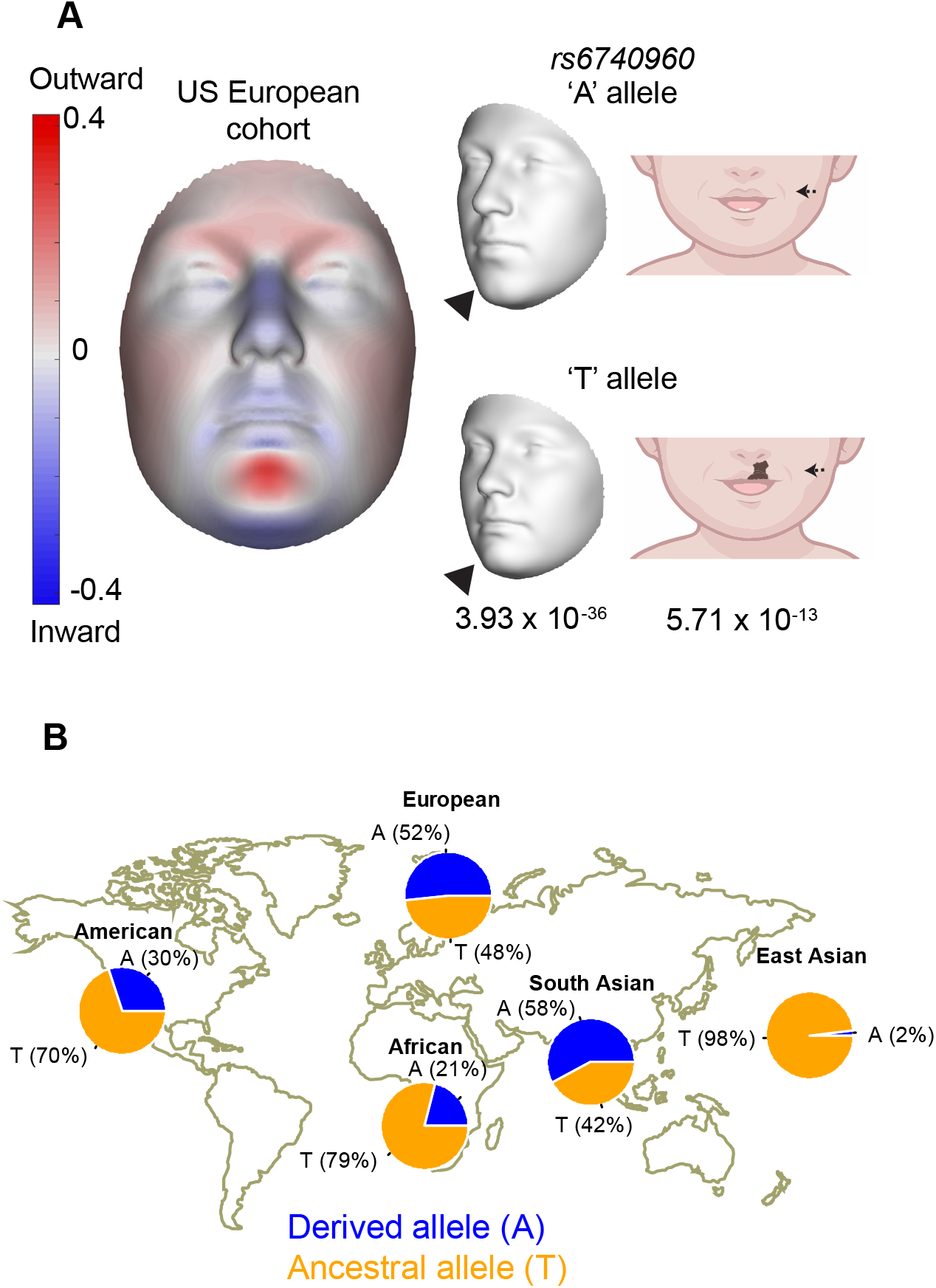
Association of *rs6740960* (A/T) with normal-range facial shape variation and its allele frequency among world populations. **(A)** GWAS of normal-range facial variation in Europeans identified *rs6740960* as a lead SNP at the 2p21 locus (Claes et al. 2018). 3D facial morphs and facial effects of the *rs6740960* are shown, with blue and red indicating a local shape depression and protrusion, respectively, due to the ‘T’ allele. Note the protrusion of the lower jaw and zygomatic regions and retrusion of the entire central midface associated with the ‘T’ allele. An independent GWAS linked *rs6740960* with susceptibility to non-syndromic cleft lip with or without palate in Europeans (Ludwig et al. 2017). **(B)** Allele frequency distribution for *rs6740960* across populations of the 1000 Genomes Project. (see also **Figure 1- Figure Supplement 1**)

We were intrigued that a common genetic variant – indeed *rs6740960* is approaching 50% allele frequency in European ancestry populations – simultaneously affects normal-range jaw shape and predisposes to clefting. To understand the prevalence of this SNP across diverse human populations, we examined allele frequencies for *rs6740960* within the 1000 Genomes Project populations (Genomes Project Consortium et al. 2015). We observed a large range of ‘A’ vs ‘T’ allele frequencies, with the “A” allele occurring at roughly 50% frequency in Europeans and South Asians (i.e. Europeans = 52%, South Asians = 58%), but at much lower frequencies in other populations (i.e. 21% in Africans and to its lowest frequency of 2% in East Asians) (**Figure 1B**). These allele frequencies were also consistent within a second human population dataset, namely the Human Genome Diversity Project (HGDP) (Bergstrom et al. 2020). Large shifts in *rs6740960* allele frequency within human populations suggest that this GWAS variant may have distinct impact on facial shape variation and disease predisposition in different populations, with potential population-specific selection events.

### *rs6740960* resides in a craniofacial enhancer with evolutionarily modulated activity

To investigate the evolutionary history of the *rs6740960* variant, we constructed sequence alignments of extant and archaic reconstructed hominin and hominid species, centered at the SNP (**Figure 2A**). These multi-species alignments revealed that the “T” allele (associated with a more protruding jaw morphology and predisposition to clefting) is the ancestral allele, shared with the Neanderthal and the chimpanzee, whereas the “A” variant at this position is a derived allele unique to humans. Interestingly, the T to A substitution at this site is associated with the gain of a putative DLX binding site (**Figure 2A**), with the DLX family transcription factors playing important roles in jaw patterning and development [reviewed in (Minoux and Rijli 2010)]. Additionally, we observed three chimp-specific mutations in the vicinity of *rs6740960*, one of which is predicted to confer gain of the ETS transcription factor family binding site (**Figure 2A**).

**Figure 2:**
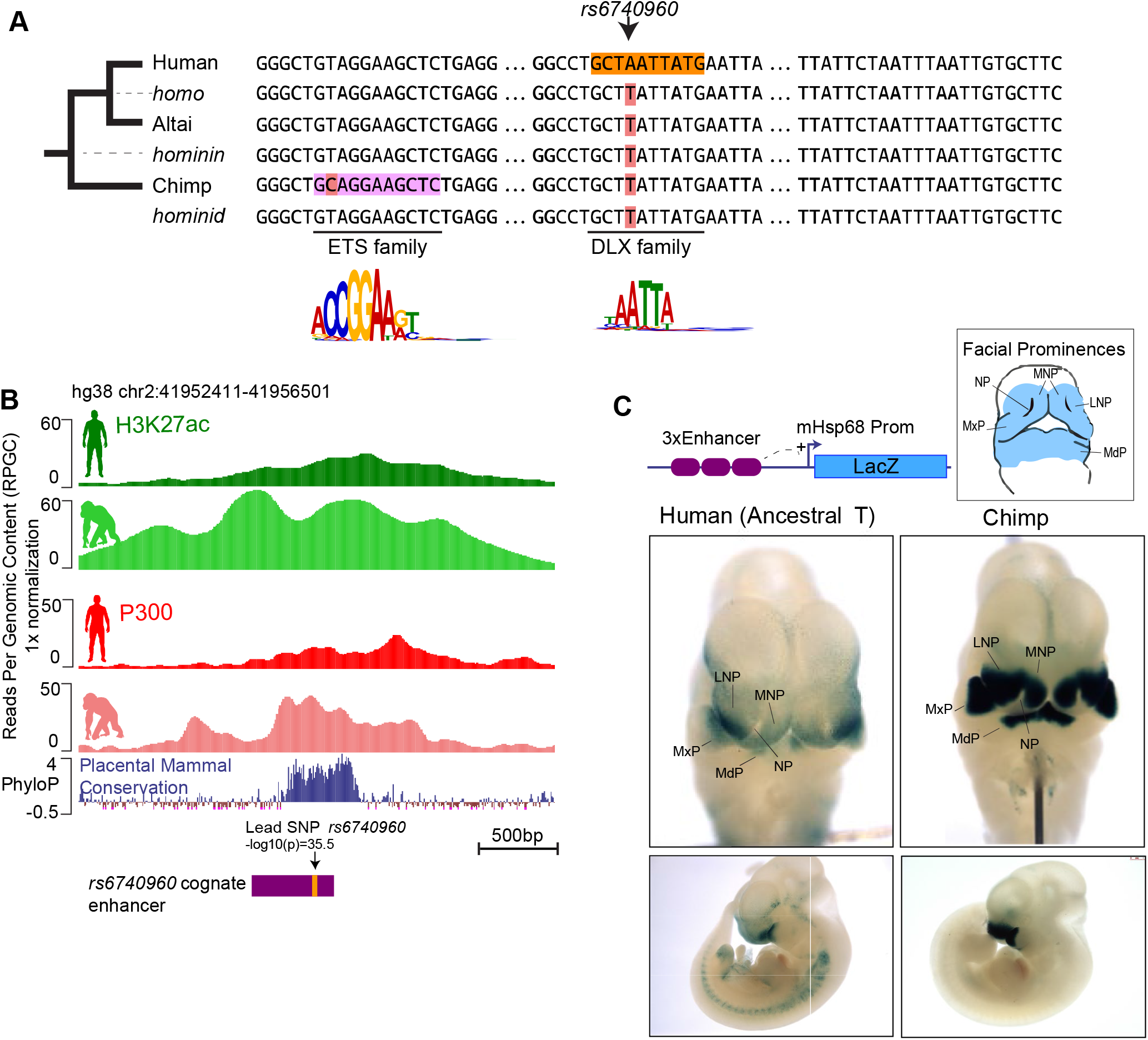
Sequence and activity changes of *rs6740960* cognate enhancer in hominids. **(A)** Multi-species abridged alignment of the genomic region surrounding *rs6740960*. Extant species are Human, Altai Neanderthal and Chimp, and ancestral reconstructed species are Homo, Hominin, and Hominid. Human-specific substitution (“A”at *rs6740960*) and chimp-specific substitution (T→C) are highlighted, along with transcription factors predicted to gain binding affinity via the substitution and their consensus sequence motifs. **(B)** Genome browser track showing the location of *rs6740960* and its overlap with H3K27ac and P300 ChIP-seq signal from human and chimpanzee CNCCs. Note higher enrichments in the chimpanzee. Region corresponding to the sequence tested in **(C)** is highlighted in purple. ChIP-seq data from (Prescott et al. 2015) **(C)** LacZ Transgenic Mouse Reporter assays performed using Human (with an ancestral “T” allele) and Chimp orthologs of the genomic region surrounding *rs6740960*. Triplicate copies of the 500bp sequence orthologs are used in the reporter vector. Both human and chimp orthologs show LacZ reporter activity restricted to the head and face prominences in E11.5 transgenic mice and overlapping upper and lower jaw primordia (Maxillary Prominence-MxP and Mandibular Prominence-MdP). The Chimp ortholog shows stronger activity within both MxP and MdP and within the Lateral and Medial Nasal Processes (LN P and MNP). NP – Nasal Pit.

These observations suggest that *rs6740960* may influence facial shape by affecting activity of a *cis-*regulatory element, and that the same element may also be modulated in a chimp-specific manner. Consistent with this hypothesis, we have previously noted that *rs6740960* resides within a genomic region marked by an active enhancer chromatin signature in *in vitro* derived human cranial neural crest cells (CNCCs) (Claes et al. 2018) (**Figure 2B**). Furthermore, enrichment of the coactivator p300 and of the active chromatin mark H3K27ac at this region is higher in the chimpanzee CNCCs as compared to human, suggesting that this non-coding element may have higher regulatory activity in the chimp (**Figure 2B**).

To test whether the genomic region spanning *rs6740960* is a *bona fide* enhancer, and if so, whether activity of this element diverged between humans and chimps, we cloned orthologous human (with an ancestral T variant at *rs6740960*) and chimpanzee regions into a LacZ reporter vector and assayed them by conventional transgenic mouse reporter assays. The resulting activity patterns were analyzed by LacZ staining of E11.5 mouse embryos. We observed that the human element displayed specific craniofacial activity in the developing upper and lower jaw primordia (mandibular and maxillary prominences), and in the lateral nasal prominence, but had little activity in the medial nasal prominence or in other tissues of the embryo (**Figure 2C**). This pattern confirms that the genomic region overlapping *rs6740960* is a *bona fide* enhancer active during craniofacial development. For simplicity, we will hereafter refer to this element as the ‘*rs6740960* cognate enhancer’. The chimp ortholog of the *rs6740960* cognate enhancer was also active in the developing face, but relative to the human element, showed both stronger and broader activity, which extended to the medial nasal prominence. Since we have examined the human ortholog with the ancestral ‘T’ variant, the stronger activity differences seen with the chimp enhancer could be due to any of the 3 additional sequence differences between human and chimp. However, the single T→C mutation that increases affinity of the chimp enhancer for the ETS family transcription factors presents the most compelling explanation of the observed activity bias. Together, our data demonstrate that *rs6740960* resides within a *bona fide* craniofacial enhancer whose activity diverged in hominids.

### The *rs6740960* cognate enhancer regulates expression of *PKDCC*

To begin probing the mechanism by which the *rs6740960* cognate enhancer can influence face shape and affect susceptibility to clefting, we first set out to identify its target gene(s). Enhancers often affect gene expression at long genomic distances and analysis of cell type-specific long-range chromatin interactions can help nominate their target genes (Jeng et al. 2019; Lopez-Isac et al. 2019). To examine long-range interactions of the *rs6740960* cognate enhancer during craniofacial development, we utilized an *in vitro* model previously developed in our laboratory, in which hESC are differentiated to CNCCs, and then further to cranial chondrocytes (Bajpai et al. 2010; Rada-Iglesias et al. 2012; Prescott et al. 2015; Long et al. 2020) (**Figure 3A**). From these cell derivatives, we profiled active chromatin by H3K27ac ChIP-seq and enhancer-promoter interactions using H3K27ac HiChIP, an assay based on principles of chromatin conformation capture, which further enriches for long-range interactions between active regulatory elements via an immunoprecipitation step with an H3K27ac antibody (Mumbach et al. 2016). In H3K27ac ChIP-seq data, we observed that the *rs6740960* cognate enhancer is comparably enriched for H3K27ac both in CNCCs and in cranial chondrocytes, although in general, there was more H3K27ac at the locus in chondrocytes (**Figure 3B**). We further confirmed commensurate H3K27ac levels at the *rs6740960* cognate enhancer in CNCCs and chondrocytes using the more quantitative ChIP-qPCR analysis (**Figure 3- Figure Supplement 1**). Surprisingly, however, in the HiChIP assay, the *rs6740960* cognate enhancer makes contacts with the promoter of the *PKDCC* gene selectively in chondrocytes, but not in CNCCs (**Figure 3B**). While we observed another significant contact between the *PKDCC* promoter and more proximal enhancers both in CNCCs and in chondrocytes, we detected significant contact between the *PKDCC* promoter and the *rs6740960* cognate enhancer only in chondrocytes. (**Figure 3B**). *PKDCC* is an attractive target gene candidate for the *rs6740960* cognate enhancer, because it encodes a secreted tyrosine kinase involved in chondrocyte differentiation, which when mutated in mice results in skeletal defects in limbs and face, including clefting (Imuta et al. 2009; Kinoshita et al. 2009; Bordoli et al. 2014).

**Figure 3:**
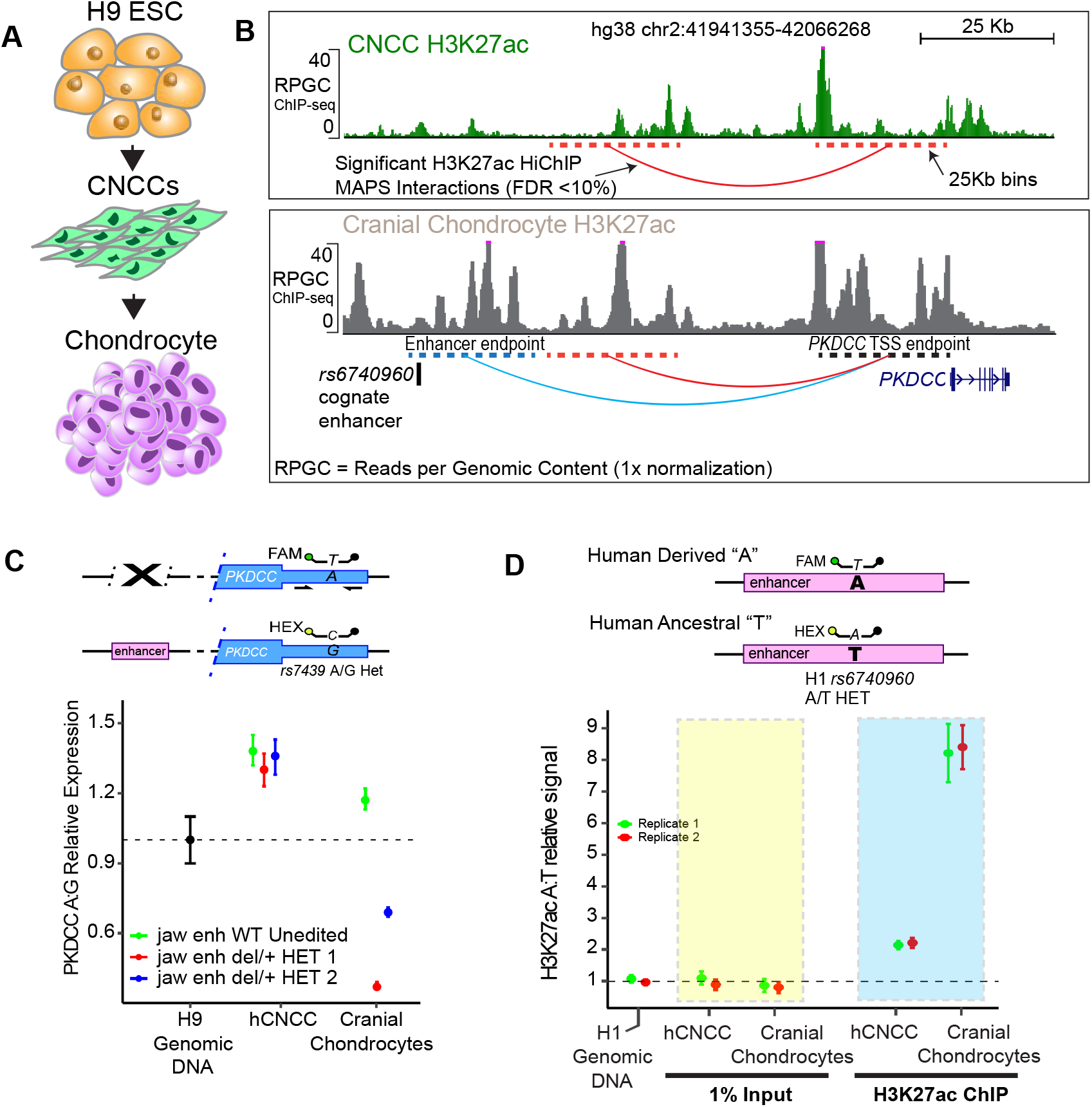
*rs6740960* cognate enhancer targets and regulates expression of *PKDCC* in cranial chondrocytes. **(A)** Schematics of the *in vitro* differentiation protocol for obtaining CNCCs and cranial chondrocytes from hESCs. **(B)** Mapping of long-range chromatin interactions by HiChI P in CNCCs and cranial chondrocytes revealed significant contacts between *rs6740960* cognate enhancer and *PKDCC* promoter selectively in chondrocytes. H3K27ac ChIP-seq genome browser tracts at the locus from CNCCs (top) and cranial chondrocytes (bottom) are shown. Significant HiChIP interactions, called at FDR < 10% and within 25Kb bins given the resolution of this dataset (dotted lines), are shown underneath H3K27ac ChIP-seq tracks. **(C)** Strong contribution of the *rs6740960* cognate enhancer to *PKDCC* expression in chondrocytes. Two independent CRISPR-Cas9 heterozygous enhancer deletion hESC lines and wildtype hESCs were di fferentiated to CNCCs and cranial chondrocytes. FAM/HEX ddPCR probes that distinguish *PKDCC* alleles based on a SNP within the 3’UTR (which was linked to the enhancer allele via phasing, see **Figure 3- Figure Supplement 3**) were used to profile allele-specific expression of *PKDCC* using RNA extracted from indicated cell lines. Relative FAM/HEX signal is shown (corresponding to ‘ A’ and ‘G’ SNP within 3’UTR, respectively). Genomic DNA from H9 wildtype cells were used to verify the FAM/HEX ratio of one. **(D)** H1 hESCs are naturally heterozygous for A/T alleles of *rs6740960*. H3K27ac-ChIP-ddPCR experiments conducted in CNCCs and cranial chondrocytes derived from H1 hESCs showed an allele-specific bias in H3K27ac signal intensity that was strongest within chondrocytes. Relative H3K27ac-ChIP-ddPCR signal at ‘ A’ vs ‘T’ allele is plotted. Note that in chondrocytes, ‘A’ allele confers an 8-fold increase in H3K27ac signal. (see also **Figure 3- Figure Supplement 1-3**)

To directly test the contribution of the *rs6740960* cognate enhancer to *PKDCC* expression, we used CRISPR/Cas9 genome editing to generate hESC lines containing a heterozygous deletion of the 1.2 kB sequence corresponding to the entire, broad p300 peak spanning the *rs6740960* cognate enhancer (**Figure 3- Figure Supplement 2**). Following differentiations of two independent heterozygous enhancer deletion lines and wild-type unedited hESCs to CNCCs and cranial chondrocytes, we profiled the allele-specific expression of *PKDCC* using droplet-digital PCR (ddPCR). In ddPCR, fluorescently labeled FAM or HEX probes designed against a heterozygous SNP (*rs7439*) within *PKDCC* 3’ UTR region permit allele-specific measurements of *PKDCC* expression in the presence or absence of the enhancer. The assay is internally controlled, since measurements of *PKDCC*’s expression with or without the enhancer are simultaneously assayed in thousands of individual droplets where each droplet receives a single template. By phasing the enhancer deletion strand with the corresponding *PKDCC* SNP allele (**Figure 3- Figure Supplement 3)**, *PKDCC*’s expression can be computed from the final number of FAM or HEX labeled droplets using Poisson statistics. These experiments showed an approximately 60% decrease in *PKDCC* expression at the allele bearing the enhancer deletion. Notably, this effect was present only within the heterozygous deletion lines but absent from wild-type cells, as expected (**Figure 3C**). In concordance with H3K27ac HiChIP interactions, the effect of the enhancer deletion was present in cranial chondrocytes, but not in CNCCs.

Both in CNCCs and in chondrocytes, we detected an allelic imbalance in *PKDCC* expression even in wild-type cells (with expression ratio from *rs7439* A vs G allele at ~1.2), indicating that other *cis-* regulatory variants at the locus can affect *PKDCC* transcript levels and potentially modulate effects of *rs6740960* on facial shape. This is consistent with the presence of many other putative enhancers at the locus, marked by H3K27ac (**Figure 3B**). Collectively, our results demonstrate that the *rs6740960* cognate enhancer targets the *PKDCC* gene and provides a strong contribution to its expression selectively in chondrocytes.

### *rs6740960* variant causes cell-type specific allelic differences in the chromatin state of its cognate enhancer

We next asked if the *rs6740960* SNP is sufficient to result in differences in chromatin state of its cognate enhancer in its native chromosomal context. We focused on H3K27ac enrichment, as this modification has been most closely correlated with enhancer activity and its changes in various biological settings (Calo and Wysocka 2013). To detect potentially subtle allelic differences in H3K27ac, we turned again to the ddPCR approach. Given that the H9 hESC line utilized in the heretofore described experiments is homozygous for the *rs6740960* SNP, we used another well-characterized hESC line, H1, which we found to be heterozygous for the *rs6740960* SNP. Following differentiation to CNCCs and cranial chondrocytes, we performed H3K27ac ChIP analyses and read out variant-specific acetylation using ddPCR assay with fluorescent probes against “A” or “T” variants at *rs6740960* (**Figure 3D**). While, as expected, there was no differences in signal in the chromatin input samples, we saw a two-fold higher H3K27ac level for the enhancer haplotype bearing the derived “A” allele in CNCCs, which further increased to approximately 8-fold difference in cranial chondrocytes (**Figure 3D**). Given that the probe-targeted SNP is the only sequence substitution within the analyzed region, we attribute observed differences in acetylation to the *rs6740960* variant, with gain of activity associated with the human-derived ‘A’ allele. Thus, the *rs6740960* drives substantial changes in enhancer chromatin state. These differences are most pronounced in chondrocytes, a cell type in which the *rs6740960* cognate enhancer contribution to *PKDCC* expression is also the highest.

### *In vivo* perturbation of *Pkdcc* dosage affects shape of jaw and palatine bones

Our molecular studies of the *rs6740960* cognate enhancer, its *cis-*mutations, and *PKDCC* expression suggest that *rs6740960* affects normal-range facial shape and predisposition to clefting by quantitatively influencing *PKDCC* dosage during cranial chondrogenesis. To directly quantify the impact of *PKDCC* dosage changes on facial morphology, we turned to mouse models. Knockout of *Pkdcc* (also known as *Vlk*) in mice results in craniofacial malformations and clefting, indicating evolutionary conservation of PKDCC craniofacial function, and nominating mice as the most relevant, experimentally accessible model for directly quantifying the impact of *Pkdcc* dosage changes on facial morphology (Imuta et al. 2009; Kinoshita et al. 2009). However, despite detectable sequence conservation of the *rs6740960* cognate enhancer in mammalian genomes (**Figure 2B**), we found that the orthologous mouse sequence lacks consistent activity in transgenic enhancer reporter assays (**Figure 4- Figure Supplement 1**), thereby limiting usefulness of the endogenous enhancer deletion experiments in mice.

In contrast to the *cis-*regulatory sequences, gene function is often deeply conserved in mammals, and this is also true for the craniofacial role of the PKDCC kinase. Given that the *rs6740960* cognate enhancer affects *PKDCC* dosage by altering its expression, we resorted to a more straightforward approach of perturbing *Pkdcc* gene dosage in mice through heterozygous coding mutations. To this end, we used previously described *Pkdcc* (*Vlk*) knockout mice (Kinoshita et al. 2009). The phenotypes reported in the *Pkdcc* −/− mice include perinatal lethality associated with respiratory and suckling defects, growth retardation, shortened limbs, delayed ossification, and craniofacial anomalies; at least some of these malformations have been attributed to the perturbed chondrogenic differentiation and maturation (Imuta et al. 2009; Kinoshita et al. 2009). Within the cranial skeleton, *Pkdcc* −/− neonates were described to have cleft palate and shortened nasal capsule and maxilla, whereas dysmorphic craniofacial phenotypes have not been reported in heterozygous *Pkdcc* +/− mice (Imuta et al. 2009; Kinoshita et al. 2009). To confirm the reported loss-of-function phenotypes and to quantitatively characterize potential effects associated with *Pkdcc* heterozygosity, we performed high-resolution micro-computed tomography (micro-CT) imaging on wild-type, *Pkdcc +/−* and *Pkdcc −/−* mice at E18.5 (**Figure 4A**). First, we confirmed that *Pkdcc −/−* embryos (n = 13) indeed displayed a shortened nasal capsule and severe clefting affecting the relevant regions of the maxilla and palatine bones as compared to wild-type embryos (**Figure 4- Figure Supplement 2, Supplementary Movies 1 and 2**).

**Figure 4:**
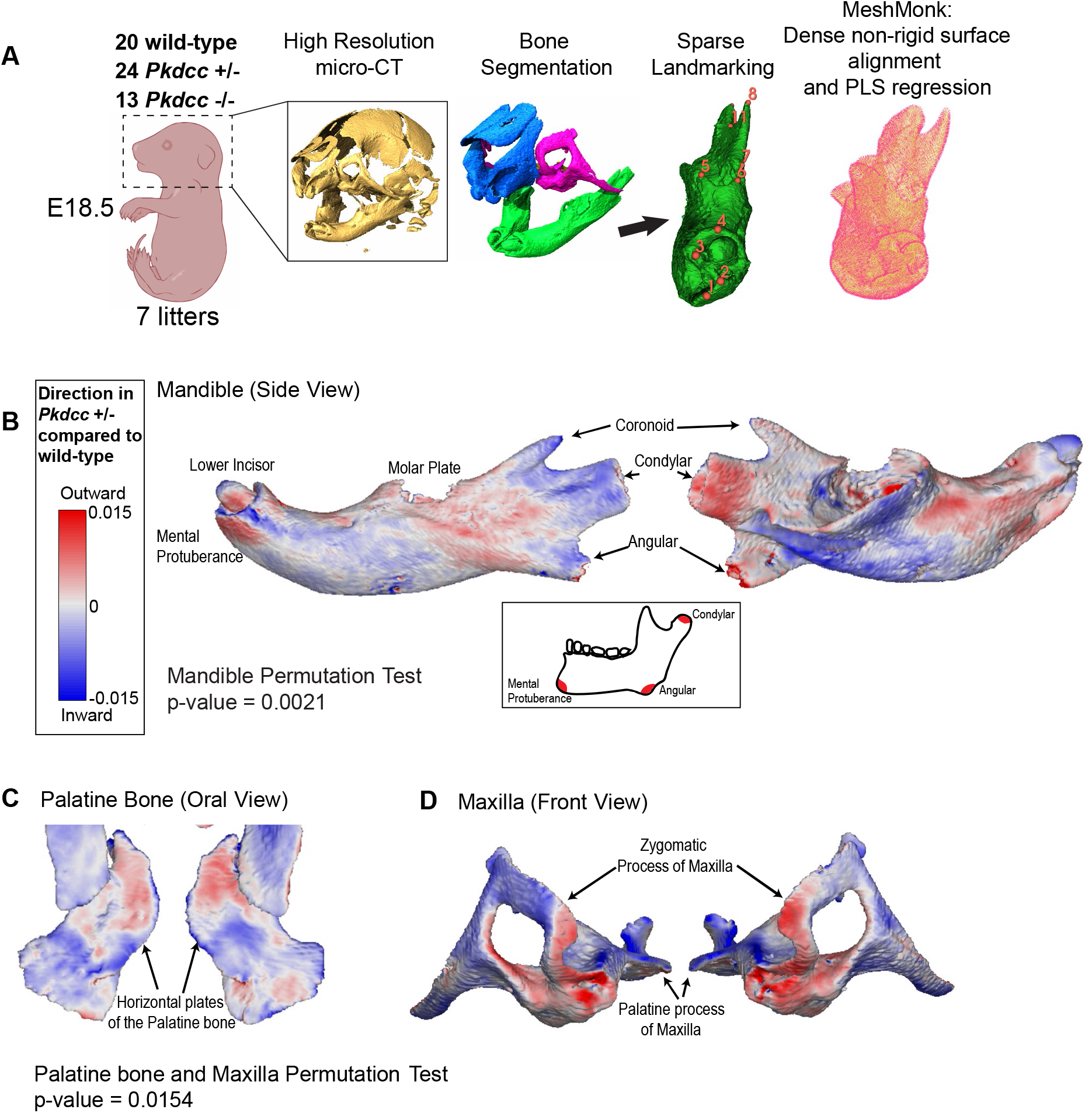
Morphogenetic assessment of *Pkdcc* dosage on cranial skeletal variation in mice. **(A)** Workflow of cranial morphometric analysis from wildtype and *Pkdcc* mutant mice. Following colony expansion, wild-type, *Pkdcc* +/− and *Pkdcc* −/− E18.5 embryos were fixed and imaged using high resolution micro-C T. Following bone segmentation and sparse landmarking, bone surfaces were non-rigidly aligned to template samples with the MeshMonk software, which facilitated dense correspondence map between all animals ‘ surfaces. Partial least squares (PLS) regression was used to analyze genotype-to-phenotype e ffects. **(B**,**C**,**D)** Bone surface variation between wild-type and *Pkdcc* +/− heterozygous mutant mice for the Mandible **(B)**, Palatine Bone **(C)**, and the Maxilla **(D)**. Blue and red coloring represent local depression and protrusion of the bone surface, respectivel y, due to 50% reduction in *Pkdcc* expression. **(B)** *Pkdcc* +/− mice exhibit overall significant shape change in the mandible (Permutation test global p-value = 0.0021). Local variation in the mandibular ramus, angular process, condylar process, and the mental protuberance contribute to the overall elongation of the mandible (See also **Supplementary Movie 3**). **(C**,**D)** *Pkdcc* +/− mice exhibit overall significant shape change in the Maxilla and Palatine bones (Permutation test global p-value = 0.015). **(C)** *Pkdcc* +/− mice have wider distance between the horizontal plates of the palatine bone, a structure that forms the posterior part of the hard palate (See also **Supplementary Movie 4**). **(D)** Significant variation in the shape of maxilla, especially within the zygomatic arches between wild-type and *Pkdcc* +/− animals. Similar analysis performed between wild-type and *Pkdcc* −/− animals are show in **Supplementary Movie 1** (mandible), and **Supplementary Movie 2** (maxilla/palatine). (see also **Figure 4-Figure Supplement 1-2**)

We next proceeded to examine the phenotypes of *Pkdcc +/−* animals. Given the modest, normal-range face shape changes associated with *rs6740960* in humans (**Figure 1A**), we also expected subtle phenotypes associated with *Pkdcc* heterozygosity in mice, requiring both a large animal sample size and sensitive morphometric analysis. In this vein, we utilized a pipeline that incorporated high-resolution micro-CT data from 20 wild-type and 24 *Pkdcc +/−* E18.5 mutant embryos spanning seven litters, manual bone segmentation, and dense landmarking of bone surfaces to capture subtle effects on bone shape variation (**Figure 4A**). To circumvent the limitations of sparsely labeled bone landmarks common to standard morphometric analysis, we used the software package MeshMonk, to densely landmark each specimen via a non-rigid registration of a template surface onto each specimen thereby allowing comparative bone surface evaluation (White et al. 2019). Using the full collection of aligned surfaces as the measured phenotype, we used partial least-squares regressions and permutation tests to compute the effect of genotype, independent of other covariates such as litter and sex, on phenotype, either at the level of complete bone shape or variation at each vertex (see **Methods**). Local regions of depression or protrusion in the shape of the bone in *Pkdcc* heterozygous mutant animals, as compared to wild-type animals, are visualized in blue and red color maps representing the regression coefficients.

Using this pipeline, we characterized the phenotypes of the mandible, maxilla, and palatine bones. Remarkably, we saw overall significant changes to the mandible shape, with strong local effects seen within the mandibular ramus and mental protuberance region, especially evident in the elongation of the angular process and mental region in *Pkdcc +/−* mutants as compared to wild-type animals, phenotypes consistent with the protruding jaw phenotype reported in GWAS (**Figure 4B, Supplementary Movie 3**). Additionally, we observed significant shape variation in the oral view of the palatine bone, especially in the distance between the symmetric horizontal plates whose eventual closure forms the base of the hard palate (**Figure 4C, Supplementary Movie 4**). A similar effect was observed, but to a lesser extent, for the palatal processes of the maxillae. The palatine and maxillary phenotypes are more subtle and distinct from the complete cleft phenotype that is present in the *Pkdcc −/−* homozygous mice. These changes can explain why alternations in *PKDCC* dosage associated with a common genetic variant in humans increase susceptibility to clefting, but are not by themselves sufficient to cause CL/P. Finally, we examined the maxilla for shape differences between genotypes. We observed no appreciable differences within the palatine process of the maxilla, but we saw significant shape changes throughout the zygomatic processes (**Figure 4D, Supplementary Movie 4**). Interestingly, these observations may potentially explain why the *rs6740960* variant shows a modest, but significant association with cheek shape in facial GWAS (**Figure 1A**). In summary, quantitative exploration of facial phenotypes in *Pkdcc +/−* heterozygous mice has allowed us to identify specific anatomical structures that are affected by decreases in *Pkdcc* gene dosage. The remarkable concordance of these phenotypes and face shape changes associated with the *rs6740960* substitution in humans provides strong support to our hypothesis that this genetic variant affects normal-range and disease-associated facial variation through regulating expression of *PKDCC*.

## Discussion

Our study provides an in-depth functional analysis of a non-coding genetic variant associated with both normal-range and disease-associated variation in a human morphological trait. Based on our collective results we propose a mechanism of *rs6740960* function. The *rs6740960* SNP resides in, and modulates activity of, a craniofacial enhancer active in the developing jaw primordia. This enhancer regulates expression of *PKDCC* in cranial chondrocytes. The *PKDCC* gene encodes a secreted tyrosine kinase, which remodels extracellular matrix during cartilage development and affects timing of proliferative chondrocyte differentiation and subsequent bone formation (Bordoli et al. 2014). Although in mice *Pkdcc/Vlk* is also important for long bone development in limbs (Imuta et al. 2009; Kinoshita et al. 2009), the craniofacial activity of the cognate *rs6740960* enhancer restricts the SNP effects on *PKDCC* dosage to the developing face. Thus, a parsimonious explanation for the association of *rs6740960* with facial variation is through the sensitivity of cranial chondrogenesis, and in turn facial skeletal development, to the *PKDCC* dosage. This may be especially pronounced during lower jaw development, where the transient Meckel’s cartilage impacts mandibular form (Svandova et al. 2020), and mandibular chondrocytes can directly transform into bone cells (Jing et al. 2015). Indeed, we demonstrated that in mice 50% reduction in *Pkdcc* dosage results in quantitative changes in shape of several mandibular regions.

Our observations can also explain why *rs6740960* is associated with nsCL/P. The reduction in *Pkdcc* dosage increased the distance between the horizontal plates of the palatine bones (and the palatal plates of the maxillae to a lesser extent) which ultimately need to fuse to form the secondary palate, but by itself, is not sufficient for clefting. This makes sense, given that the frequency of the risk-conferring *rs6740960* T allele in the European ancestry population (even in a homozygous setting) is two orders of magnitude higher than the frequency of orofacial clefting in the same population. Thus, by definition, the risk variant must work additively with other variants acting on genes affecting palate development, either at the same locus or at different loci, and/or with environmental factors. These include prenatal exposure to alcohol, nicotine, viruses, maternal diabetes, and methyl donor deficiency, all of which are linked to an increased prevalence of orofacial clefts (Martinelli et al. 2020). Interestingly, unaffected relatives of individuals with nsCL/P show distinctive facial features, which have been quantitatively characterized to define a subclinical ‘endophenotype’ that may reflect a heightened susceptibility to clefting (Indencleef et al. 2021; Weinberg 2022). Subsequent GWAS analysis revealed loci associated with nsCL/P endophenotype in a healthy, unselected population (Indencleef et al. 2021). Notably, one of the genome-wide significant lead SNPs from this analysis, *rs4952552*, maps to the *PKDCC* locus, suggesting that the impact of *rs6740960* on clefting risk is likely modulated by the presence of other common variants acting on *PKDCC*.

It is not entirely clear what are the embryological origins of the maxillary and palatine changes observed in *Pkdcc* heterozygous mutants, particularly given the intramembranous origin of these bones. One possibility is that these changes are a secondary consequence of effects taking place in the adjacent endochondral cranial base. Another possibility worth future exploration is that in addition to chondrocytes, mesenchymal tissues forming the bony structures of the palate are also sensitive to *Pkdcc* dosage. Consistent with this possibility, Kinoshita et al. (2009) showed that *Pkdcc* is expressed in the CNCC-derived mesenchyme populating the palatal shelves. Regardless, the quantitative phenotypes we describe in *Pkdcc* heterozygous mice are consistent with an increased susceptibility of clefting in Europeans carrying the risk-conferring *rs6740960* allele.

The activity of the *rs6740960* cognate enhancer diverged in hominids, with both stronger and expanded facial activity domain observed in the chimpanzee, suggesting that this element may have contributed to the divergence in craniofacial morphology between hominid species. Expansion of the activity domain observed in chimp spans facial regions that will ultimately form a nose. Thus, in the chimp, genetic variation in this enhancer may conceivably affect nasal shape, in addition to the effects on the jaw shape observed in humans. Our analysis also revealed drastic variation in *rs6740960* allele frequency across modern human populations, with Europeans and South Asians showing high heterozygosity for this SNP, and East Asians exhibiting prevalence of the ancestral ‘T’ allele. What could be the potential adaptive advantages for the increased frequency of the human-derived allele in European and South Asian populations? Our analysis suggests that the variation in jaw phenotype associated with the *rs6740960* was partly driven by variation in the angular process of the mandible. The angular process, or mandibular angle, attaches to the masseter muscle which generates the majority of bite-force during mastication (Sella-Tunis et al. 2018). Could the derived “A” allele facilitate masticatory adaptations influenced by dietary changes along population boundaries? Was the increase in the derived allele frequency associated with the elevated susceptibility to clefting associated with the ancestral allele in European population? And if so, are East Asians buffered from these effects by other, compensatory genetic variants? Or, alternatively, could the advantageous benefit be purely an aesthetic one, with a less or more protruding jaw appearance affecting sexual selection differently in distinct human populations and cultures? While we do not have answers to these questions at present, our results merit further consideration in the context of human facial adaptations and evolution.

## Methods

### Cell culture and differentiation of hESCs to CNCCs and cranial chondrocytes

Human Embryonic Stem Cell (hESC) lines (H9 and H1 lines; WiCell WA09 and WA01) were cultured and differentiation to Cranial Neural Crest Cells (CNCCs) using protocol previously described in (Prescott et al. 2015). Briefly, hESCs were plated onto tissue-culture 6-well plates pre-coated with Matrigel (Corning 356231) using mTESR-1 medium (StemCell Technologies 85850) and grown for 5-8 days with daily medium replacement. Cells were passaged 1:6 using RELESR (StemCell Technologies 05872) for detachment of hESCs. For differentiation of hESCs to CNCCs, large hESC colonies were detached using 2mg/mL Collagenase IV (Fisher 17104019) resuspended in KnockOut DMEM (Gibco 10829018), and incubation for 30 min – 1hr at 37ºC. Intact cell clumps were washed in PBS and plated into Neural Crest Differentiation medium (NDM) in 10cm Petri Dishes incubated at 37ºC. Media was change daily until Day 4, changing plates every day. On Day 4, neuroectoderm spheres were left undisturbed until Day 7 to promote attachment to dish. From Day 7 – 11 CNCCs emerged from attached neural spheres, and NDM media was changed daily without disturbing attached spheres and CNCCs. To transition CNCCs to passage 1, neural spheres were removed by gentle aspiration, the plate was washed with PBS, and the remaining CNCCs were detached using 50% Accutase diluted in PBS. After incubation at 37ºC for 1 minute, CNCC were disaggregated with gentle pipetting. CNCCs were plated onto fibronectin coated 6-well tissue-culture treated plates with Neural Crest Maintenance media (NMM), left to attached for 15 minutes at 37ºC, and medium containing Accutase was replaced with fresh pre-warmed NMM. Medium was replaced daily without tilting plates. Cells were split 1:3 every 2 days, using 50% accutase, and the same procedure of allowing CNCCs to attach to the plate for 15 minutes at 37ºC, and immediately replacing accutase containing medium with fresh pre-warmed NMM. One day after passage 3, cells were transitioned to NMM + BMP2 and ChIRON medium (NMM+BC).

Passage 4 CNCCs were harvested for experiments, or differentiated further to cranial chondrocytes. For cranial chondrocytes differentiation, NMM+BC medium was replaced with Chondrocyte Differentiation Medium (CDM) without TGFβ3 supplement, with an intermediate PBS wash between media change (day 0). Medium was changed at day 1 and day 4 with CDM supplemented with TGFβ3, and cells were harvested at day 5. To disassociate single-cell chondrocytes from the extracellular matrix, we used protocol described in (Makki et al. 2017). Briefly, cells were washed with PBS, and incubated with digestion medium (1mg/mL Pronase (Roche 10165921001), 1mg/mL Collagenase B (Roche 11088815001), 4U/mL Hyaluronidase (Sigma H3506), resuspended in KnockOut DMEM) for 1 hour at 37ºC, with agitation every 15 minutes. Digestion medium was aspirated after centrifugation (5 min, 100 r.c.f), and cell pellet was washed twice with PBS, with centrifugation between each wash.

#### Neural Crest Differentiation Medium (NDM)

1:1 Neurobasal Medium (Thermo Scientific 21103049) and DMEM F-12 medium (GE Healthcare SH30271.01), 0.5X Gem21 NeuroPlex supplement (B-27) (Gemini 400-160), 0.5X N2 NeuroPlex supplement (Gemini 400-163), 20ng/mL bFGF (PeproTech 100-18B), 20ng/mL EGF (PeproTech AF-100-15), 5ug/mL bovine insulin (Gemini 700-112P), 1X Glutamax supplement (Thermo Fisher 35050061), 1X Antibiotic-Antimycotic (Gibco 15240062).

#### Neural Crest Maintenance Medium (NMM)

1:1 Neurobasal Medium (Thermo Fisher 21103049) and DMEM F-12 medium (GE Healthcare SH30271.01), 1mg/mL BSA (Gemini 700-104P), 0.5X Gem21 NeuroPlex supplement (B-27) (Gemini 400-160), 0.5X N2 Neuroplex supplement (Gemini 400-163), 1mg/mL BSA (Gemini 700-104P), 20ng/mL bFGF (PeproTech 100-18B), 20ng/mL EGF (PeproTech AF-100-15), 1X Glutamax supplement (Thermo Scientific 35050061), 1X Antibiotic-Antimycotic (Gibco 15240062).

#### NMM+ BMP2/ChIRON

NMM medium supplemented with 50pg/mL BMP2 (PeproTech 120-02) and 3uM CHIR-99021 (Selleck Chemicals S2924).

#### Chondrocyte Differentiation Medium (CDM)

DMEM High Glucose Medium (Cytiva SH30243.01), 5% FBS, 1X ITS+ Premix (Corning 354352), 1X Sodium Pyruvate (Gibco 11360070), 50ug/mL L-Ascorbic Acid (Sigma-Aldrich A4403), 0.1uM Dexamethasone (Alfa Aesar A17590), TGFβ3 (PeproTech 10036E), 1X Antibiotic-Antimycotic (Gibco 15240062).

### H3K27ac HiChIP and ChIP

CNCCs and Chondrocytes were detached as singled cells and resuspended in BSA and Serum free NMM or CDM respectively (1 million cells/mL), fixed in 1% formaldehyde with end-over-end rotation at room temperature for 10 minutes, quenched with 0.2M glycine solution for 5 minutes, and pelleted by centrifugation (5 min, 1350 r.c.f, 4ºC). Cell pellet was washed twice with cold PBS, with centrifugation between washes, and cell pellet was flash frozen in liquid nitrogen and stored at −80°C.

HiChIP experiments were conducted following protocol described in (Mumbach et al. 2016) using MboI digestion, and ChIP experiments were performed as described in (Long et al. 2020). In both experiments, we used Active Motif Rabbit Polyclonal H3K27ac ChIP-grade antibody (Cat No 39133). HiChIP experiments were performed on eight biological replicates of CNCC and chondrocyte differentiations, and both HiChIP and 1% Input libraries were sequenced on Illumina NovoSeq 6000 (2 × 100bp) platform resulting in 244 million non-duplicate reads. HiChIP data was processed using HiC-Pro pipeline with 25Kb resolution (Servant et al. 2015), the recommended resolution of this data prescribed by HiC-Res (Marchal et al. 2020). HiC-Pro was used as a preprocessing workflow to aligns reads to the hg38 genome, and to save only valid read pairs (i.e. from *cis* long and short range interactions, and intra-chromosomal interactions). Significant 3D interactions (10% False Discovery Rate) were called using MAPS (Juric et al. 2019).

To call significant 3D interactions, MAPS requires precomputed 1-dimension (1D) ChIP peaks, ideally called from independent ChIP-seq experiments. For this purpose, H3K27ac ChIP-seq peaks for CNCCs were identified using published Human H3K27ac ChIP-seq data from (Prescott et al. 2015). To call ChIP-seq peaks in cranial chondrocytes, we performed H3K27ac ChIP-seq experiments from three additional chondrocyte differentiations. ChIP-seq data were aligned to the hg38 genome using BWA MEM aligner (Li and Durbin 2009), and broad peaks were identified using macs3 (Zhang et al. 2008). Finally, for visualizing comparable H3K27ac ChIP signal for CNCC and chondrocytes in Figure 3B, we performed H3K27ac ChIP-seq on a single hESC → CNCC → chondrocyte differentiation.

### LacZ transgenic mouse experiments

Enhancer elements were cloned in triplicate copy into a LacZ reporter plasmid immediately upstream of a Hsp68 promoter and LacZ-P2A-tdTomato fusion gene, all surrounded by core insulator elements. Orthologs of the *rs6740960* cognate enhancer from Human, Chimp, and Mouse (each 500bp in length) were amplified from genomic DNA extracted from Human H9, Chimp C0818, and Mouse E14 ESC cell lines. H9 is homozygous for the “A” (derived) allele of *rs6740960*. To create a human homolog bearing the “T” ancestral allele, we utilized PCR primer bearing this allele and PCR fragment ligation to recreate a mutagenized sequence. Mouse transgenesis experiments, conducted by Cyagen, were performed as described in (Samuels et al. 2020). Briefly, reporter plasmids were linearized, injected into fertilized mouse oocytes, where they randomly integrated into the genome, oocytes were implanted into recipient females, and allowed to develop to E11.5 embryonic stage. Embryos were harvested, genotyped for positive reporter integration, and stained with X-gal to reveal the anatomical structures where the reporter transgene was expressed. We required at least 3 LacZ positive embryos with consistent LacZ staining profiles for expression domains to be consider reproducible.

### Generation of enhancer knock-out cell lines using CRISPR/Cas9

Deletion of the *rs6740960* cognate enhancer in H9 hESCs was accomplished with CRISPR/Cas9 genome editing using the CRISPR RNA “GTGGGGATTGCGCTAACTCA” synthesized by IDT and homology-direct repair (HDR) template bearing the 1.2Kb enhancer deletion. HDR template was cloned into an AAV2 production plasmid (Addgene 32395), using left and right homology arms cloned from H9 genomic DNA (Left Arm Primers GTGCAGAGCTGTCTG / ATGTTTTACTAGCTCCTGCTAT; Right Arm Primers AGGGAGGATGGGAAGGA / GACAGGCAAGAGACTGACATA). AAV HDR donor plasmid, and Capsid 2 and AD5 helper plasmids, were co-transfected into HEK-293FT cells (Invitrogen R70007), and after 48 hours, high-titer virions were purified by Iodixanol ultracentrifugation using a modified protocol provided in (Martin et al. 2019).

H9 hESCs were cultured in mTESR-1 medium supplemented with 10mM ROCKi (Y27632; StemCell Technology 72304) for 24hrs, washed with PBS, and brought to single-cell pellet with Accutase treatment for 5 min at 37ºC and gentle centrifugation (100 r.c.f; 5 min). 1 million hESCs were resuspended with 100uL Lonza Amaxa P3 nucleofection solution (Lonza V4XP-3024), the annealed, pre-complexed crRNA:tracrRNA guide-RNA duplex (3uL of 50uM gDNA duplex), and 19ug Alt-R S.p. HiFi Cas9 v3 Nuclease (IDT 1081060) (2.5:1 Cas9:gRNA molar ratio). Cells were electroporated using the CA-137 setting on the Lonza 4D Amaxa Nucleofector, immediately plated onto precoated matrigel plates with 100K MOI AAV2 HDR, and cultured with mTESR1 + 10mM ROCKi for 96 hours. Edited single-cells were passaged sparsely (600 cells per well of 6-well plate), and allowed to grow for 7 days. Colonies were manually picked, expanded, and genomic DNA was isolated using a Quick-Extract Buffer (Lucigen GE09050). Successful heterozygous deletion lines were identified using the PCR primers CCCCCACCCATCAGTCATTC and GGTTGTGCCTCATAGTGCCT and Q5 Polymerase (NEB M0491S). Edited lines showed a 2.4kB band corresponding to the deletion allele in contrast to the 3.6kB unedited, wildtype band. To phase the enhancer deletion allele with *PKDCC* heterozygous SNP *rs7439*, we used the PCR primers CAGCAGCCAGAGATCCTTTAG and AAGGAAAGCCTCCACTTCTTT to amplify a 12.7kB region encompassing the enhancer and *rs72792594*, the closest pre-phased SNP obtained from H9 10X whole-genome sequencing data (Long et al. 2020).

### Droplet Digital experiments for *rs6740960* H3K27ac signal and PKDCC expression

Droplet digital PCR (ddPCR) experiments were performed using protocols for the PCR master mix, droplet generator and droplet reader provided by BioRad. To assay allele-specific H327ac-ChIP signal at the *rs6740960* cognate enhancer, CNCCs and cranial chondrocytes cells differentiated from H1 hESCs were fixed and ChIP’ed as described above, and ChIP DNA (50X dilution) was used as input in the ddPCR experiments. To validate the that fluorescently labeled probes designed against biallelic SNPS were unbiased in their binding affinities, H1 genomic DNA was used as a quality control to ensure 50:50 ratio of droplets. To assay allele-specific expression of *PKDCC, rs6740960* cognate enhancer heterozygous deletion H9 lines and wild-type H9 cells were differentiated to CNCCs and chondrocytes. Total RNA was extracted using TRIzol Reagent according to manufacturer’s protocol (Thermo Scientific 15596026), and 2ug of total RNA was DNase I treated and reverse-transcribed to cDNA using random Hexamer oligos according to Themo SuperScript IV VILO kit (Thermo Scientific 11766050). cDNA was diluted 40X for used as input DNA in RT-ddPCR experiments.

IDT Locked Nucleic Acid (LNA) FAM and HEX fluorescently-labeled probes were designed to distinguish biased H3K27ac ChIP signal at the A/T alleles of *rs6740960* in H1 cell derivatives (56-FAM/ATT+CATAA+T+T+AGCA+GGCCATG/3IABkFQ/ and /5HEX/ATT+CATAA+T+A+AGCA+GGCCATG/3IABkFQ/). Forward and reverse PCR primers to amplify the 119bp region surrounding *rs6740960* were CTTTCTTCTGTGAGCTGTAGCA and TCCACGCTGGGTTTAGTTATTT. Similarly, probes were designed to assay the allele-specific expression of PKDCC in *rs6740960* cognate enhancer heterozygous deletion H9 cell derivatives. Specifically, FAM/HEX probes designed against heterozygous SNP *rs7439* (A/G) located within S*PKDCC*’s 3’UTR (/56-FAM/CGG+G+A+G+A+TGC/3IABkFQ/ and /5HEX/CGG+G+G+GATGC/3IABkFQ/). This 88bp 3’UTR region was amplified with TCGTGGAGTGTTCTCTCA and AGCCTCCCCAGGTAC PCR primers.

BioRad QuantSoft Analysis Pro v1.0 software was used to analyze ddPCR experiments. QuantSoft uses Poisson statistics to estimate the number of FAM positive and HEX positive droplets in the starting reaction. These ratios were plotted in R to understand the allelic bias in H3K27ac ChIP and allele-specific *PKDCC* expression.

### Micro-CT of E18.5 PKDCC transgenic mice

*Pkdcc* knock out mice were created by (Kinoshita et al. 2009) by replacing the first exon with a Venus - SV40 polyA tag. Cryogenic preserved two-cell oocytes were obtained from RIKEN BioResource Research Center (RBRC057275) and rederived at Stanford University’s Transgenic, Knockout, and Tumor Model Center (TKTC). Founder animals were genotyped using a triplicate primer set (GGCATGGACGAGCTGTACAA, GGCTCAGGCTACACTAAGGC, CTGGGTTCTGCACCTAGCTG), that resulted in a single 332bp amplicon for wildtype animals, a single 594bp amplicon for homozygous-tagged animals, and dual bands for heterozygous animals. Mice were housed at the Animal Veterinary Center at Stanford University. All procedures involving animal care and use were conducted under a pre-approved protocol by the Administrative Panel on Laboratory Animals Care at Stanford University (APLAC Protocol No. 30364).

Mouse embryos were collected at E18.5 developmental stage, and fixed in 4% Paraformaldehyde for 7 days at 4ºC. Embryos were imaged on Bruker Skyscan 1276 micro-CT imager using the following settings (15um pixel size, 2K resolution, 0.25mm Al filter, ~750ms exposure, 360º scanning, rotation step 0.5 degree, 100% partial width, 2 frame averaging). Bruker NRecon software was used for 3D reconstruction using the settings smoothing = 2, ring artifacts correction= 20, beam hardening = 0%, Gaussian Smoothening Kernel. Volume-of-interest (VOI) were isolated for the head region of each animal using Bruker DataViewer software. Mandible, maxilla, and palatine bones were segmented and roughly landmarked for each VOI using Amira 3D 2021.1 software. We used landmarks recommended by (Ho et al. 2015). Segmented bone surfaces were exported as Wavefront OBJ format and landmarks were exported as fiducial CSV formats for morphometry analysis.

### Morphometry analysis for E18.5 embryos

Morphometric differences in cranial bone surfaces of wildtype, *Pkdcc* +/−, and *Pkdcc* −/− E18.5 embryos were analyzed in Matlab 2021b. The MeshMonk toolbox (White et al. 2019) was developed for spatially dense phenotyping of 3D surface meshes of anatomical structures and has been used extensively in studies of craniofacial variation in humans (Claes et al. 2018; Liu et al. 2021; White et al. 2021). MeshMonk performs a non-rigid registration of a template mesh onto each target mesh. Thereafter each target sample is represented in terms of the topology of the template mesh. The vertices of each registered surface are spatially-dense analogues of traditional anthropometric landmarks. In this study, the mandible, maxilla, and palatine bone surfaces from one representative wildtype animal was chosen as the template sample and the corresponding structures in each sample were registered with this template.

Further statistical analysis was performed using in-house software developed by authors HM and PC in MATLAB 2021b. All landmark configurations were aligned to their mean and scaled to unit size by generalized Procrustes analysis. Dimensionality of landmark variation was reduced by projection onto the principal components (PCs) explaining 96% of variation. The partial effect of genotype (independent of litter and sex) on the shape was tested with partial least-squares regressions and permutation test (10,000 permutations) on the partial effect size (partial R^2^) as described in (Shrimpton et al. 2014). The categorical covariates sex and litter were each coded as k-1 binary ‘dummy’ variables, where k is the number of categories. To test for effects at the level of the whole structure, principal component scores were normalized to have unit variance along each PC, and the partial effect size is the variance explained in these normalized PCs. To test for effects at each vertex the principal component projection was reversed and partial R^2^ was computed for each vertex. The regression coefficients represent the change in the shape of the bone between wild-type and *Pkdcc* mutant animals are visualized as color maps showing change in the direction perpendicular to the surface at each vertex (blue=inward; red=outward).

## Data Availability

H3K27ac ChIP-seq and HiChIP-seq sequencing data generated in this study are available from the Gene Expression Omnibus (GEO Accession Number: GSE212234).

## Materials Availability

Edited human ESC cell lines (H9) and other research reagents generated in this study are available to academic researchers by contacting the corresponding author.

## Supplementary Materials

**Figure 1- Figure Supplement 1:**
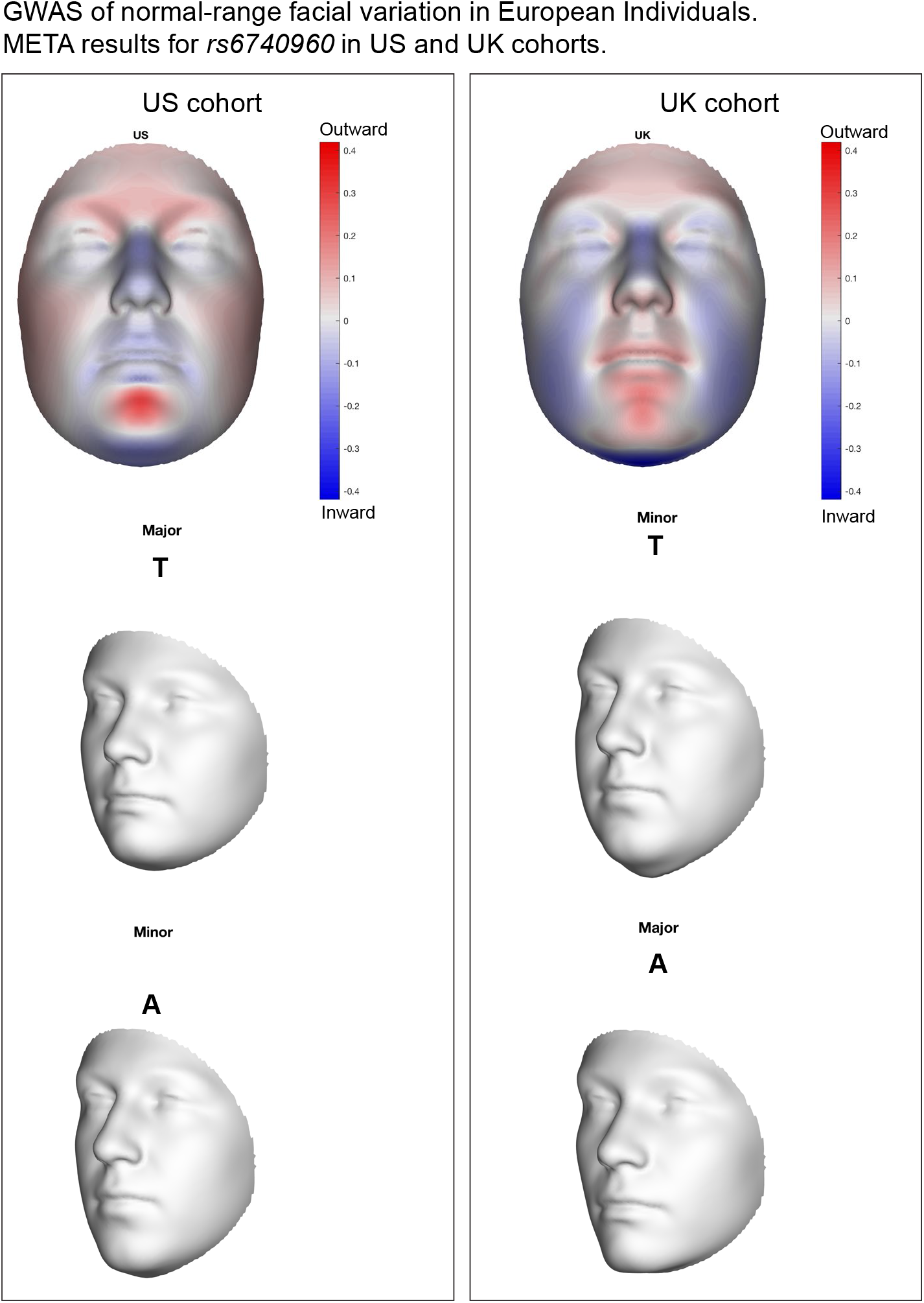
Facial effects for the *rs6740960* (A/T) SNP implicated in normal-range facial variation within the United Kingdom and United States of America European study cohorts of (White et al. 2021). Blue/red coloring indicates depression/protrusion local variation due to the effect of the “T” allele. Gray 3D face morphs show the averaged effects of the “A” or “T” alleles within either the UK or US cohorts from (White et al. 2021). Note that the “minor” and “major” allele designations are reversed amongst these two cohorts due to *rs6740960’s* approximate 50% heterozygosity in European populations.

**Figure 3- Figure Supplement 1:**
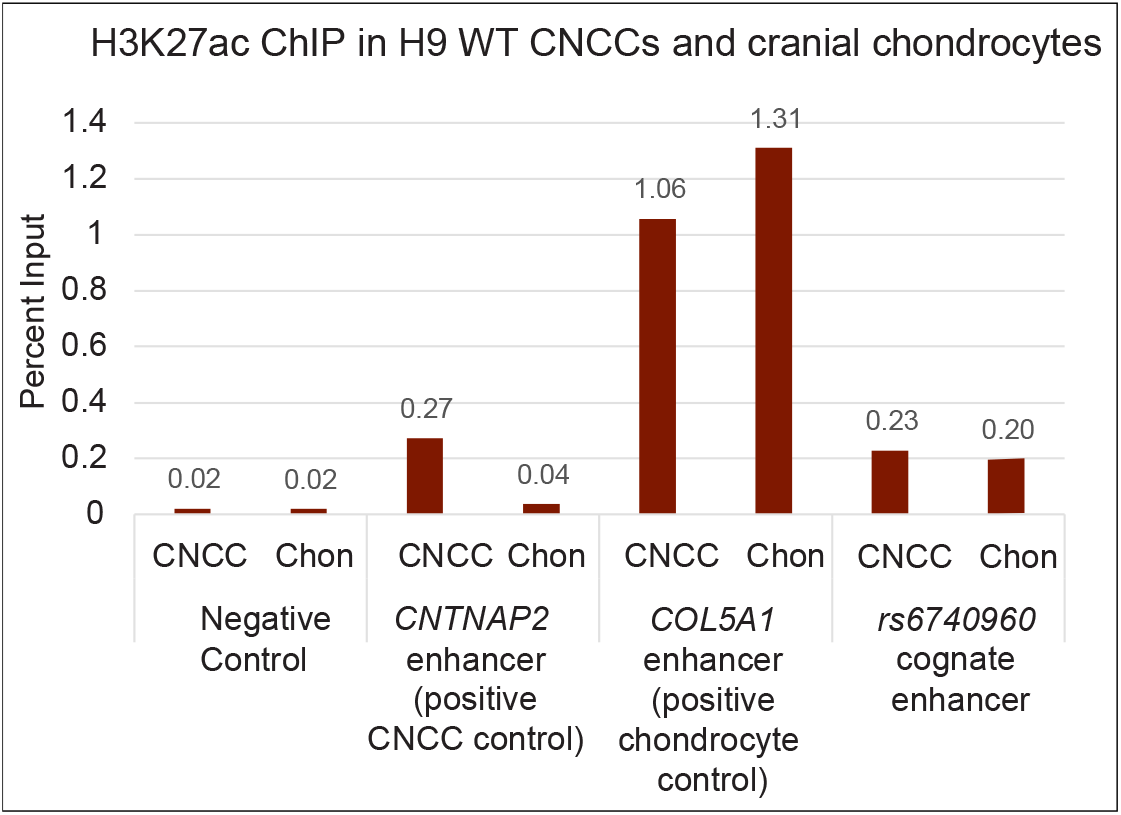
H3K27ac ChIP-qPCR data from H9 hESC derived CNCCs and cranial chondrocyte cells, as shown in **Figure 3A** and **3B**. Primers are designed for an intergenic region devoid of active chromatin marks (negative control), a CNCC-enriched enhancer adjacent to the gene *CNTNAP2*, a chondrocyte-enriched enhancer adjacent to the *COL5A1* gene, and the *rs6740960* cognate enhancer. H3K27ac-qPCR data indicate similar deposition of this chromatin modification mark at the *rs6740960* cognate enhancer in CNCCs and cranial chondrocytes.

**Figure 3- Figure Supplement 2:**
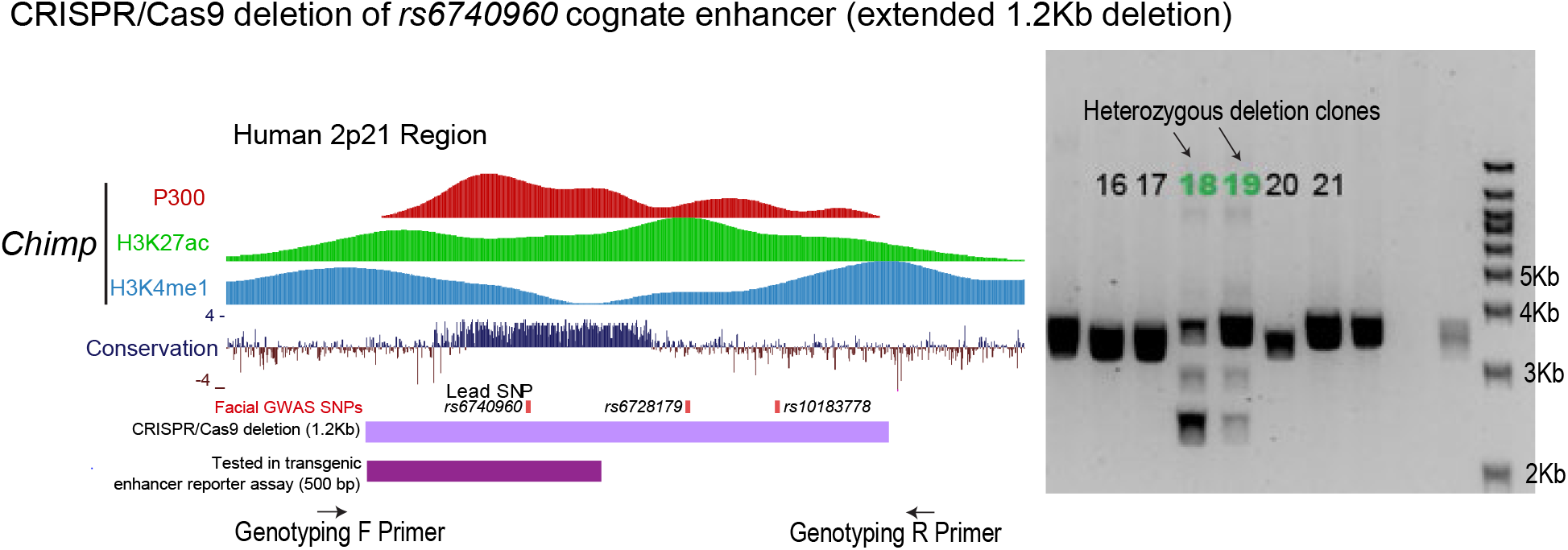
Generation of a 1.2Kb deletion at the *rs6740960* cognate enhancer using CRISPR/Cas9 genome editing of H9 hESCs. Genome browser tracks depicting P300, H3K27ac, and K3K4me1 ChIP-seq data from Chimpanzee C0818 cell-line collected in (Prescott et al. 2015). The P300 profile guided our selection of the 1.2Kb enhancer region for deletion. Face-shape associated GWAS SNPs from (White et al. 2021) are noted. Genotyping PCR results of clonal hESC lines indicate two heterozygous enhancer deletion clones (#18 and 19).

**Figure 3- Figure Supplement 3:**
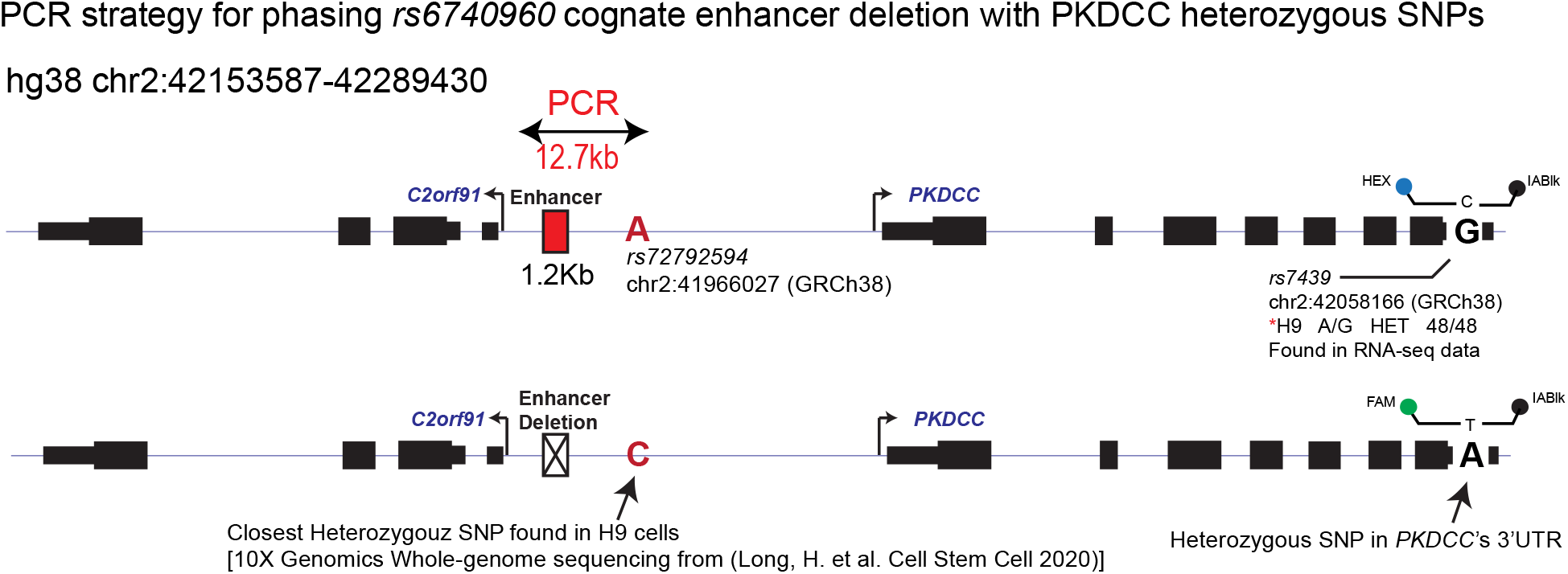
PCR strategy for phasing the 1.2Kb enhancer deletion with *PKDCC* 3’UTR heterozygous SNP (*rs7439*). PCR primers were used to amplify a 12.7Kb region spanning the enhancer deletion with the closest pre-phased heterozygous SNP (*rs72792594*) within H9 hESCs, as identified by 10X Genomics whole-genome sequencing data from (Long et al. 2020). Sanger sequencing data of the resulting PCR amplicons support the enhancer deletion allele occurs with the “C” allele of *rs72792594*, and therefore with the “A” allele of *rs7439*.

**Figure 4- Figure Supplement 1:**
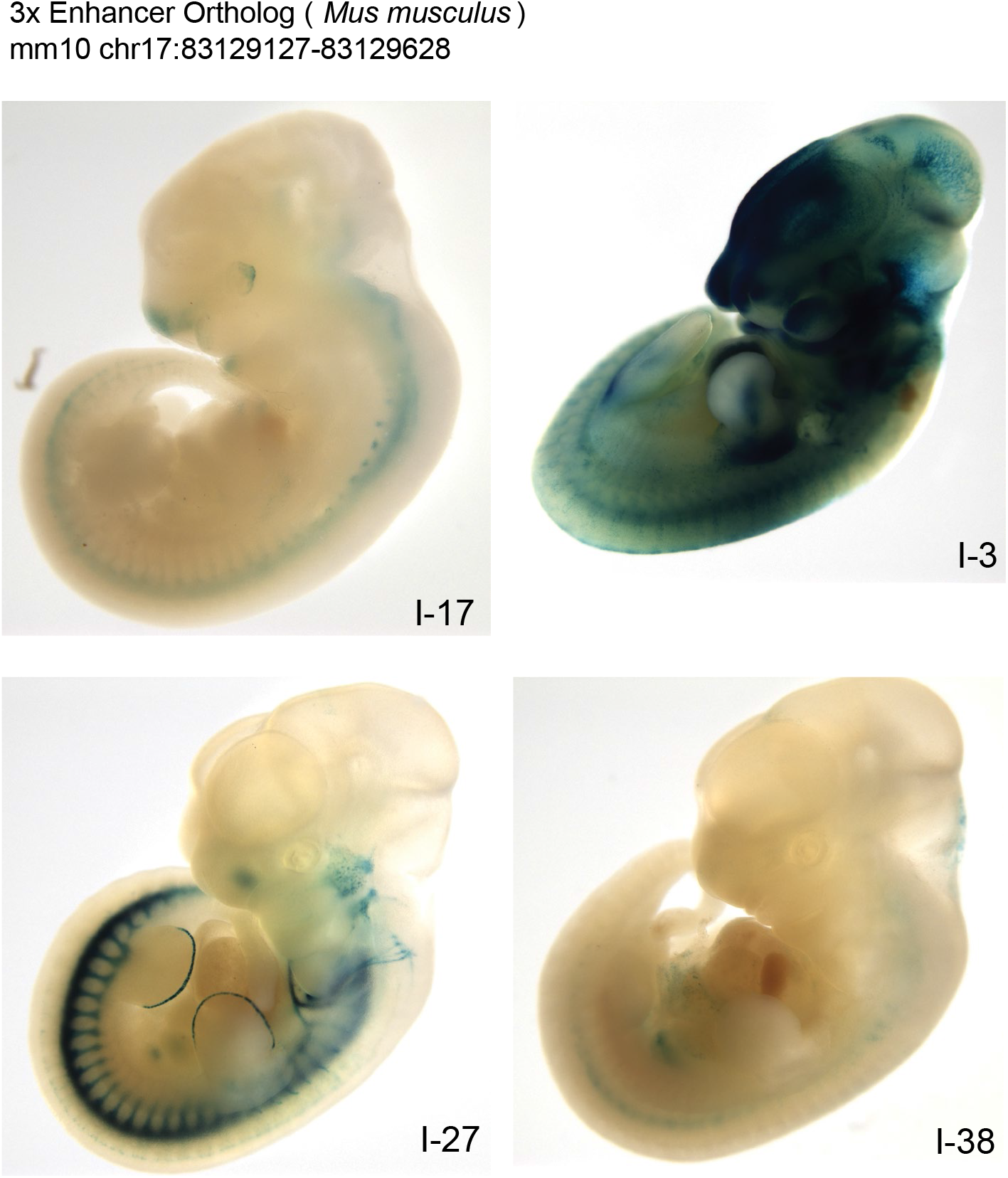
LacZ transgenic mouse reporter assay for the 500bp mouse ortholog of the *rs6740960* cognate enhancer. Similar to the human and chimp orthologs (**Figure 2C**), triplicate copies of the mouse ortholog was cloned into the LacZ reporter construct and tested. The mouse enhancer sequence does not show consistent LacZ reporter activity across animals that are PCR positive for reporter transgene integration.

**Figure 4- Figure Supplement 2:**
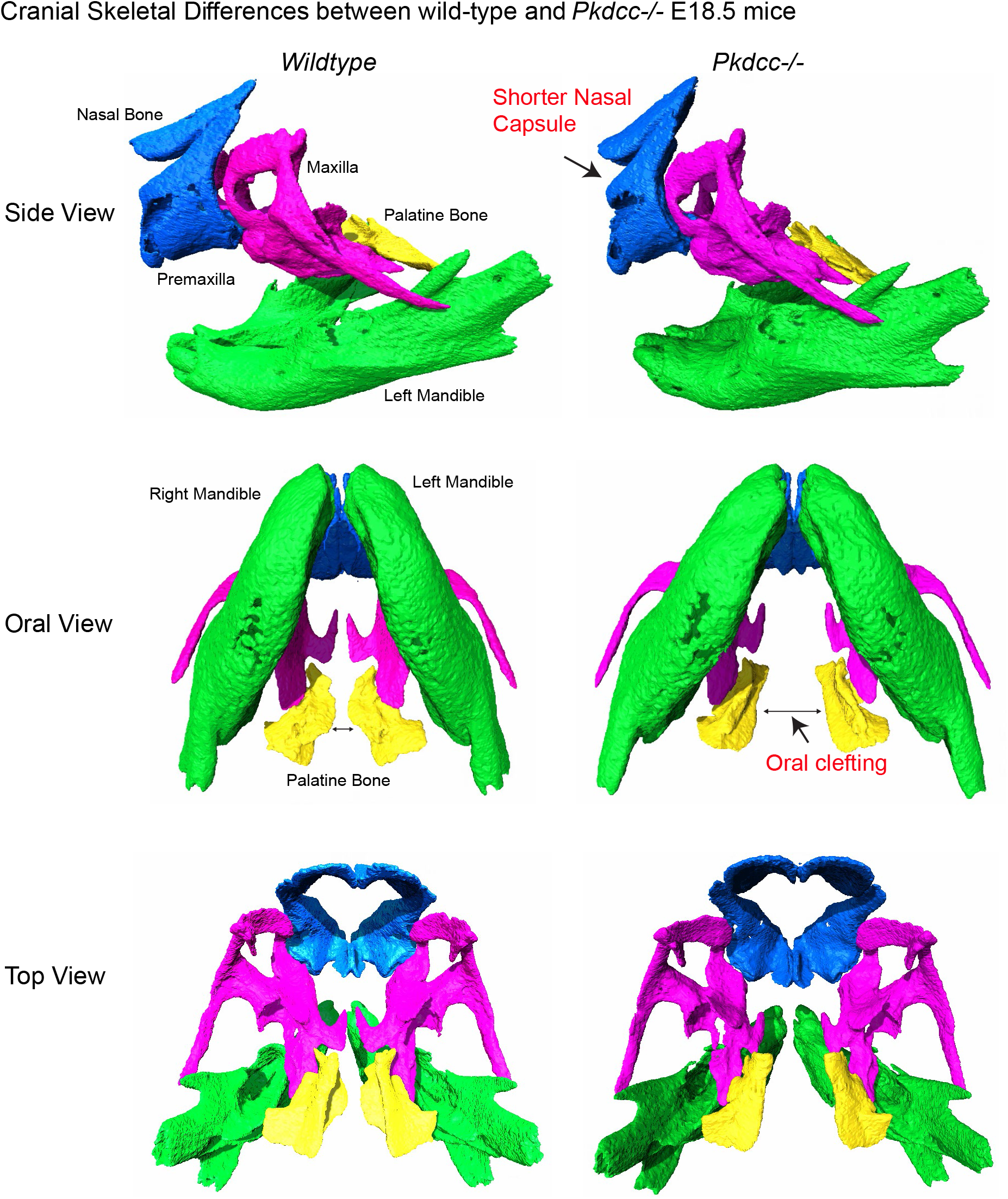
Micro-CT images demonstrating cranial skeletal differences between wildtype and *Pkdcc* −/− mutant E18.5 mouse embryos. As reported in previous studies (Imuta et al. 2009; Kinoshita et al. 2009), *Pkdcc* −/− animals have shorter nasal capsule, and oral clefting phenotypes compared to wild-type animals. Cranial bone colors are nasal and premaxilla bones (blue), maxilla (pink), left and right mandibles (green), and palatine bone (yellow).

**Supplementary Movie 1:** Left mandible shape effect changes between averaged wildtype and *Pkdcc* −/− E18.5 mouse embryos. The movie progresses from the phenotype in wildtype embryos to *Pkdcc* −/− embryos and back to wildtype embryos. **(A)** medial side view, and **(B)** lateral side view. Movies depict significant shape variation in the mandibular ramus and mental regions in *Pkdcc* −/− embryos.

**Supplementary Movie 2:** Maxilla and palatine bone shape effect changes between averaged wildtype and *Pkdcc* −/− E18.5 mouse embryos. The movie progresses from the phenotype in wildtype embryos to PKDCC −/− embryos and back to wildtype embryos. **(A)** oral view, **(B)** top view, and **(C)** front view. Movies depict a widened distance between the symmetric palatine process of the maxilla and the horizontal plates of the palatine bone in *Pkdcc* +/− embryos.

**Supplementary Movie 3:** Left mandible shape effect changes between averaged wildtype and *Pkdcc* +/− E18.5 mouse embryos. The movie progresses from the phenotype in wildtype embryos to *Pkdcc* +/− embryos and back to wildtype embryos. **(A)** medial side view, and **(B)** lateral side view. Movies depict elongation in the angular process and shape variation within the mental protuberance region in *Pkdcc* +/− embryos.

**Supplementary Movie 4:** Maxilla and palatine bone shape effect changes between averaged wildtype and *Pkdcc* +/− E18.5 mouse embryos. The movie progresses from the phenotype in wildtype embryos to PKDCC +/− embryos and back to wildtype embryos. **(A)** oral view, **(B)** top view, and **(C)** front view. Movies depict a widened distance between the symmetric horizontal plates of the palatine bone and shape variation within the zygomatic arches in *Pkdcc* +/− embryos.

## Acknowledgements

We thank members of the Wysocka lab for helpful discussions, comments, and advice. We thank Naz Koska for help with mouse husbandry, Larissa Sambel for help with cell culture and ddPCR experiments, and Drs. Heather Szabo Rogers and Sahin Naqvi for comments on the manuscript.

*Pkdcc/Vlk* targeted *Mus musculus* strain (No. RBRC057275), originating from (Kinoshita et al. 2009), was obtained from RIKEN BioResource Research Center (Acc No. CDB0434K and website http://www2.clst.riken.jp/arg/mutant%20mice%20list.html)

This research was supported by the NIH grant R01 DE027023 and funding from the Howard Hughes Medical Institute. J.W. was supported by a Lorry Lokey endowed professorship, and a Stinehart Reed award. J.M. was supported by an NSF Postdoctoral Research Fellowship in Biology (Award No. 1711847).

